# Genomic and environmental factors shape gill microbiome activity in an Amazonian teleost holobiont

**DOI:** 10.1101/2022.05.27.493806

**Authors:** François-Étienne Sylvain, Nicolas Leroux, Éric Normandeau, Aleicia Holland, Sidki Bouslama, Pierre-Luc Mercier, Adalberto Luis Val, Nicolas Derome

## Abstract

Fish microbiomes provide functions critical for their host’s survival in contrasting environments. These communities are sensitive to a range of environmental-specific factors (i.e. physicochemical parameters, free-living bacterioplankton) and host-specific factors (i.e. host genetic background). The relative contribution of these genomic and environmental factors shaping Amazonian fish microbiomes is still unknown. Here, we investigated this topic by analyzing the gill microbiomes of 240 wild flag cichlids (*Mesonauta festivus*) from four different populations (genetic clusters) distributed in 12 sites of two contrasting water types (ion-poor/acidic black water and ion-rich/circumneutral white water). The transcriptionally active gill microbiomes were characterized by a 16S rRNA metabarcoding approach carried on RNA extractions. They were analyzed in light of comprehensive datasets from the hosts genetic background (Genotyping-By-Sequencing), the bacterioplanktonic pool of bacteria (16S rRNA) and a set of 34 environmental parameters. Results show that the transcriptional activity of gill microbiome samples was significantly different between the genetic clusters and between water types. However, they suggest that the contribution of the host’s genetic background was relatively weak in comparison to the environment-related factors in structuring the relative abundance of different gill microbiome transcripts. This result was also confirmed by a mixed-effects modeling analysis, which suggested that the dissimilarity between the transcriptional activity of bacterioplanktonic communities possessed the best explicative power regarding the dissimilarity between gill microbiomes transcripts, while pairwise fixation indexes (F_ST_) from the hosts’ genetic data only had a weak explicative power. We discuss these results in terms of microbiome assembly processes and flag cichlid fish ecology.

**Importance:** Host-associated microbial communities respond to a range of factors specific to the host physiology, genetic backgrounds and life history. However, these communities also show different degrees of sensitivity to environment-dependant factors such as abiotic physico-chemical parameters and ecological interactions. The relative importance of host-versus environment-associated factors in shaping teleost microbiomes is still understudied and is paramount for their conservation and aquaculture. Here, we studied the relative importance of host- and environment-associated factors structuring teleost microbiomes using gill samples from a wild Amazonian teleost model (*Mesonauta festivus*) sampled in contrasting habitats along a 1500 km section of the Amazonian basin, thus ensuring high genetic diversity. Results showed that the contribution of the host’s genetic background was weak compared to environment-related bacterioplanktonic communities in shaping gill microbiomes, thereby suggesting that our understanding of teleost microbiome assembly could benefit from further studies focused on the ecological interplay between host-associated and free-living communities.

## 1. Introduction

The Amazon River basin is one of the Earth’s most diversified aquatic ecosystems. This region shows substantial spatial heterogeneity due to the existence of dramatic hydrochemical and ecological gradients that impose physiological constraints upon its aquatic communities (Duarte *et al*. 2016; Henderson and Crampton 1997; Junk *et al*. 1983; Petry *et al*. 2003; Rodriguez and Lewis 1997; Saint-Paul *et al*. 2000). Essentially, two distinct water types, “white” and “black” waters, result from contrasting physicochemical profiles and are characterizing the Amazon River basin (Sioli 1984). White water environments are eutrophic (nutrient- and ion-rich with high productivity), have a circumneutral pH and typically show a high turbidity (Gaillardet *et al*. 1997; Dal Pont *et al*. 2017; Sioli 1984; Val and Almeida-Val 1995). In contrast, black water environments are oligotrophic (nutrient- and ion-poor with low productivity) and contain a high concentration of dissolved organic carbon (DOC) enriched in organic acids such as humic substances. The relatively high amounts of humic DOC acidify black water environments (pH 3.0-5.0), making them physiologically challenging for local fish, especially regarding the homeostasis of ionoregulatory processes (Duarte *et al*. 2016).

Previous studies have examined the effects of these water types on various aspects of Amazonian fish ecology, especially on their phylogeography (Cooke *et al*. 2014; Pires *et al*. 2018; Leroux *et al*. 2022), migratory patterns (Hermann *et al*. 2016), evolutive strategies (Borghezan *et al*. 2021) and global community diversity (Bogotá-Gregory *et al*. 2020). However, up to now only one investigation has studied the effects of these water types on native fish microbiomes (Sylvain *et al*. 2019). Host-associated microbial communities can often be seen as an extension of the host genome by providing functions critical for the host survival in changing environments (Henry *et al*. 2021). Together with the host, they constitute a holobiont, a term coined by Margulis & Fester(1991), referring to the co-dependence between the host and its microbial symbionts. Just like the host fish transcriptome (Araújo *et al*. 2017), fish microbial communities are very sensitive to slight variations in environmental physicochemical parameters (Sylvain *et al*. 2016), and thus probably show water-type specific profiles.

In addition to environment-dependant factors such as water types, Amazonian fish microbiomes could also be influenced by their hosts’ genetic backgrounds, which are known to affect the taxonomic profiles of fish microbial communities in a variety of fish clades from other ecosystems, such as brook charr, Atlantic salmon, stickleback, elasmobranchs, whitefish and Amazonian piranhas (Boutin *et al*. 2014; Smith *et al*. 2015; Sevellec *et al*. 2018; Webster *et al*. 2018; Doane *et al*. 2020; Sylvain *et al*. 2021). However, the covariation between fish genotypes and the composition of their associated microbial communities is not always clear, as some studies suggest that environmental influences overcome host genomic effects on fish microbiome, rendering the detection of the latter difficult (Riiser *et al*. 2020; Bledsoe *et al*. 2018).

The flag cichlid (*Mesonauta festivus*) has recently emerged as an interesting model to study the factors shaping Amazonian fish microbiomes (Sylvain *et al*. 2019, 2020). The flag cichlid is a small (15 cm max), sedentary, detritivore and gregarious species that is abundantly found on the margins of almost all Amazonian lakes and rivers (Pires *et al*. 2015). They are naturally found in both black and white water environments, and thus enable the study of the effect of environment-specific factors on fish microbiomes. Furthermore, a recent analysis of the flag cichlid phylogeography has shown that water types had a low predictive power on the genetic structuration observed in this species, which was better explained by a strong influence of past vicariant events (Leroux *et al*. 2022). In their study, Leroux *et al*. (2022) showed that flag cichlids were divided into four genetically differentiated populations (referred to as “genetic clusters” in this manuscript), two of them encompassing ecosystems of both black and white water (Suppl. Fig. 1). Overall, this simple system composed of four main genetic clusters and two water types constitutes an interesting model to study the relative contribution of genomic and environmental factors shaping Amazonian fish microbiomes.

Here, our study aimed to characterize how the gill microbiome of wild flag cichlids changes according to the environmental water type and the host genetic cluster. We analyzed a dataset characterizing the gill microbiome from 240 flag cichlids distributed throughout 12 sampling sites of black and white water environments. These individuals were also used in the phylogenomic investigation of Leroux *et al*. (2022) and thus represent the four genetic clusters previously identified in that study. We chose to study the gill microbiome, rather than the gut or the skin mucus microbiomes, because the gill is the organ mostly responsible for ionoregulatory processes associated to the fish physiological response to different water types (reviewed in Morris *et al*. 2021). We used a 16S rRNA metabarcoding approach based on RNA extractions to characterize the transcriptionally “active” part of the gill microbiome. We quantified to what extent microbiome samples cluster according to their water type and their fish host’s genetic cluster. Then, we identified bacterial biomarkers specific to different water types and genetic clusters. Finally, we modeled the contribution of both environment-specific factors (i.e. free-living bacterioplankton transcriptional activity and environmental physico-chemical parameters) and host-specific factors (i.e. list of Single Nucleotide Polymorphisms used to define the genetic clusters) on gill microbiomes beta-diversity.

## 2. Results

After sequencing, merging and quality filtering, 5,102,333 reads (mean of 21,620 reads/sample, N = 236 samples) were obtained for the gill microbiome dataset. Gill microbiomes of flag cichlids showed an important relative abundance of Proteobacteria (mean of 65 ± 4 %), Bacteroidetes (26 ± 4 %) and Firmicutes (7 ± 2 %) transcripts (Fig. 2). Gill microbiomes from black water fish were significantly enriched in Firmicutes (12 ± 5 % in black water, 4 ± 1 % in white water, Student T test pvalue = 0.06), while gill microbiomes from white water specimens were significantly enriched in Bacteroidetes (34 ± 4 % in white water, 15 ± 2 % in black water, Student T test pvalue = 0.03).

**Figure 1:**
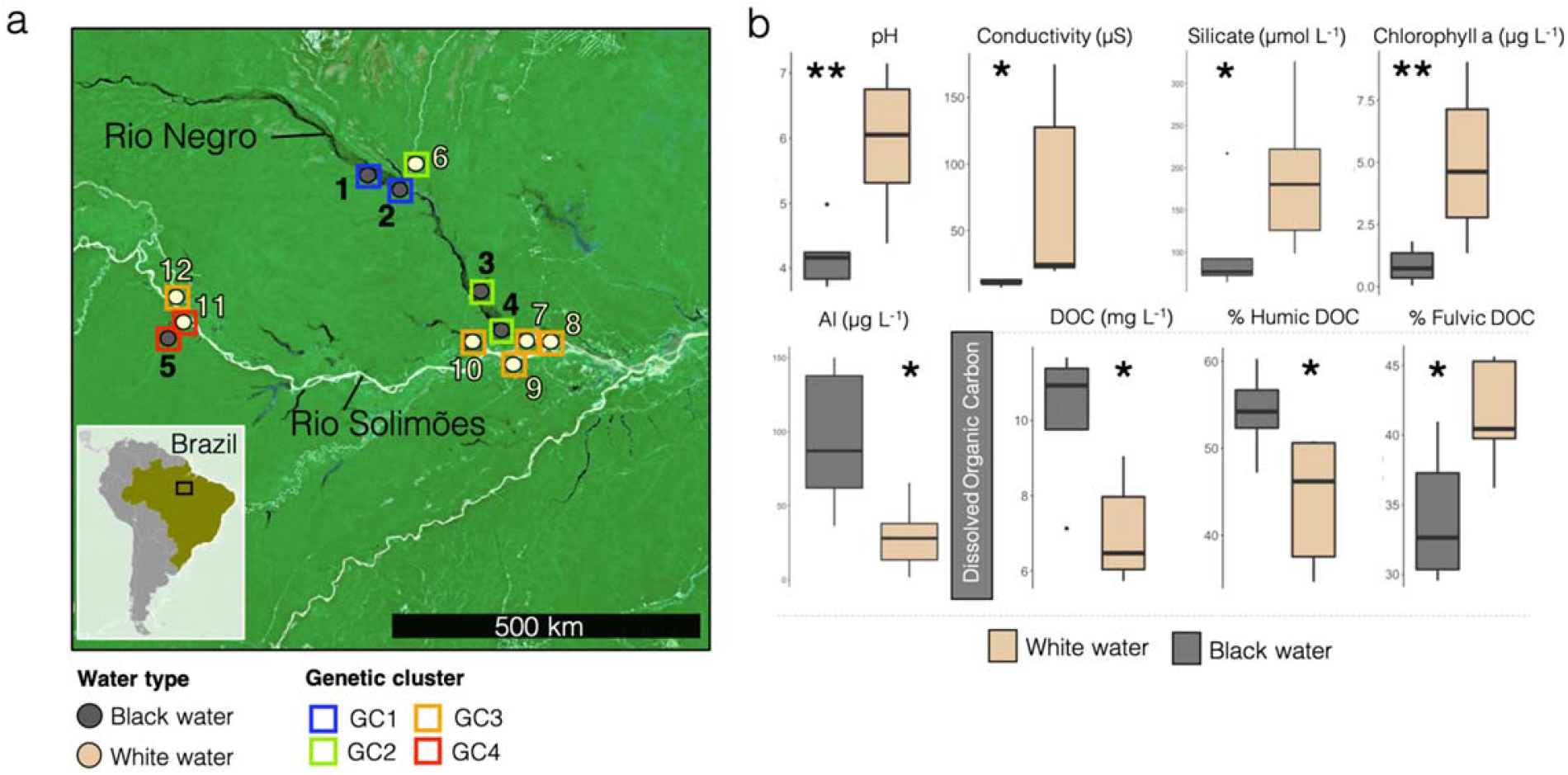
(a) Map of the 12 sampling sites including information on the water type and the genetic cluster of fish found at each site. “GC” stands for “Genetic cluster”. (b) Observed values of the environmental parameters commonly used to differentiate black from white water. “Al” refers to dissolved aluminum, “DOC” refers to dissolved organic carbon.

**Figure 2:**
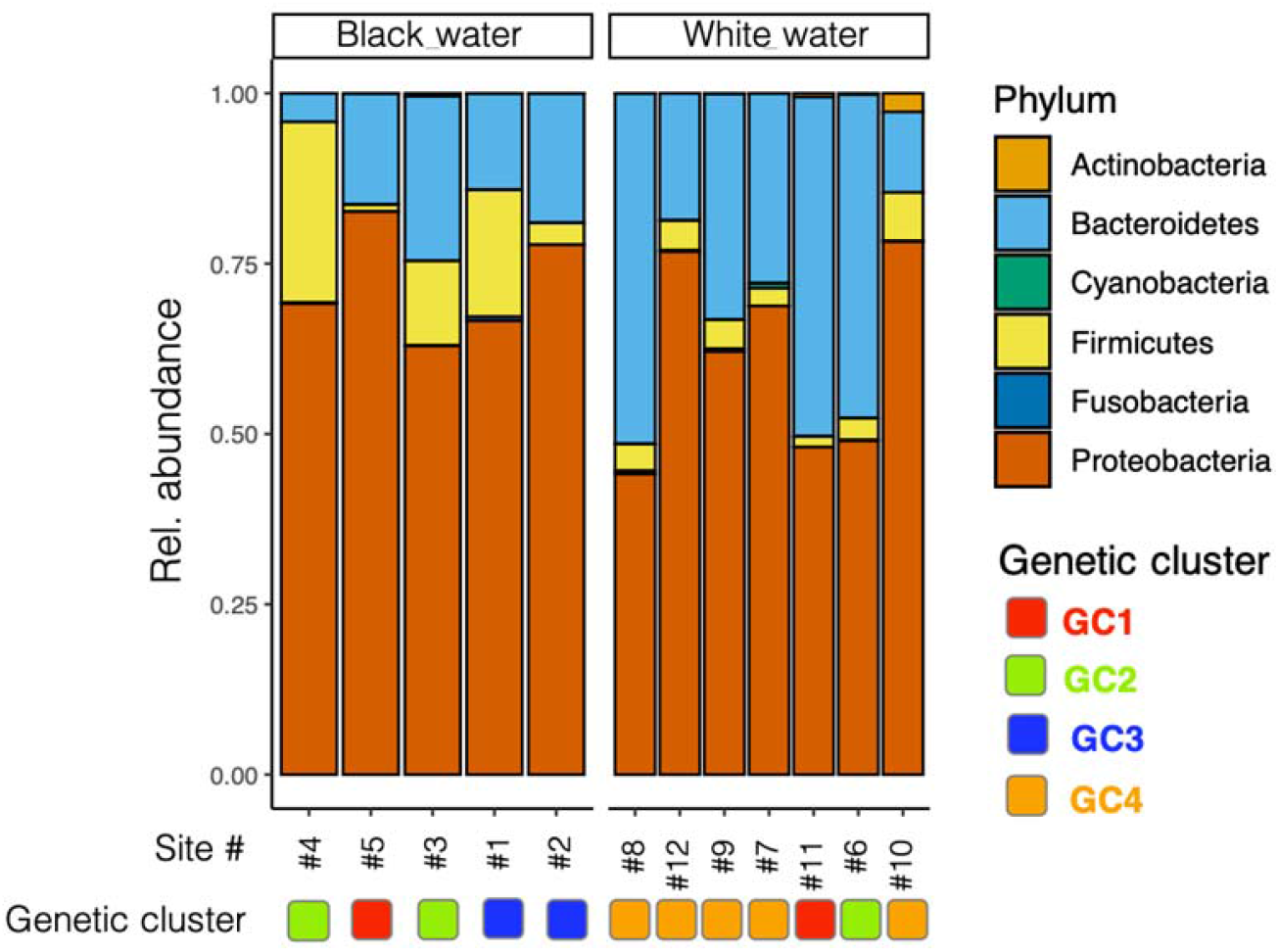
Relative abundance of the phyla detected in gill microbiome samples, according to the water type and the genetic cluster of the fish found at each sampling site. “GC” stands for “Genetic cluster”.

### 2.1 Gill microbiomes from different water types and host genetic clusters

The PCoAs conducted on Fig. 3a show that, to a certain extent, gill microbiome samples clustered according to their genetic cluster of origin. The first two axes of these PCoAs represented 21.1% and 23.3% of the variance. The results from PERMANOVAs also suggested that the gill microbiome transcriptional activity significantly differed according the genetic cluster of the host fish, both in black (F = 7.49, R^2^ = 13%, df res = 97, pval < 0.001) and in white water (F = 10.76, R^2^ = 14%, df res = 113, pval < 0.001) environments. In addition, the PCoAs from Fig. 3b suggested that gill microbiomes samples clustered according to their water type of origin. This result was obtained for the genetic clusters GC2 and GC4 for which we found samples in both black and white water types (this was not the case for the genetic clusters GC1 and GC4, thus they were excluded from this analysis). The first two axes of the two PCoAs in Fig. 3b represented 37.0% and 36% of the variance. PERMANOVAs confirmed that the gill microbiome transcriptional activity significantly differed according the water type of the sampling environment, both in genetic clusters GC2 (F = 14.30, R^2^ = 20%, df res = 57, pval < 0.001) and GC4 (F = 9.33, R^2^ = 20%, df res = 37, pval < 0.001).

**Figure 3:**
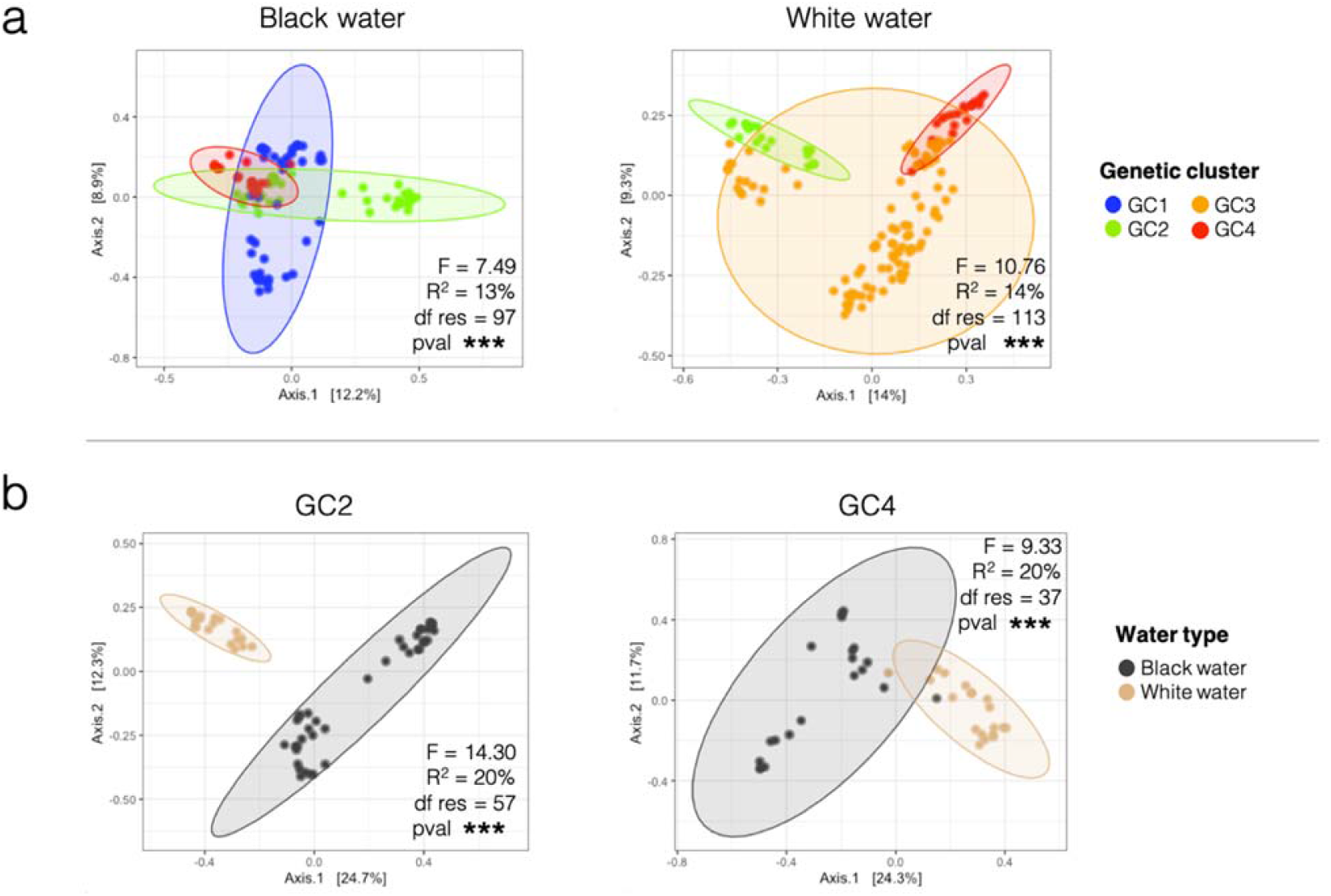
Principal coordinates analyses (PCoA) of gill microbiome samples, based on Bray-Curtis distances. Results of permutational analyses of variance (PERMANOVA) based on Bray-Curtis distances are shown at the bottom or top right corner of each plot. “df res” stands for “residual degrees of freedom”. “pval ***” stands for “pvalue < 0.001”. In (a) samples are colored according to their genetic cluster of origin. “GC” stands for “Genetic cluster”. In (b) samples are colored according to their water type of origin. The genetic clusters GC1 and GC3 are not shown in (b) as they were not found in both water types.

Then, we identified bacterial biomarkers (at the ASV level) specific to the four genetic clusters (Fig. 4a,b) or the two water types (Fig. 4c,d). We identified 90 bacterial biomarkers specific to one of the genetic clusters (23 ASVs for GC1, 9 for GC2, 29 for GC3, 29 for GC4). Their taxonomic assignations mostly represented members of the classes Alphaproteobacteria, Betaproteobacteria, Flavobacteriia and Clostridia (Fig. 4b). Biomarkers from the Clostridia class were mostly associated to GC1 and GC2, those from the Flavobacteriia and Gammaproteobacteria classes were enriched in fish from GC3, while those from the Alphaproteobacteria and Betaproteobacteria class were found in fish from all genetic clusters (GC1, 2, 3, 4) but were mostly abundant in GC4. In average, these biomarkers represented a relative abundance of 13.8 % of the global gill microbiome (11.5% in GP1, 7.0% in GP2, 10.0% in GP3, 26.5% in GP4).

**Figure 4:**
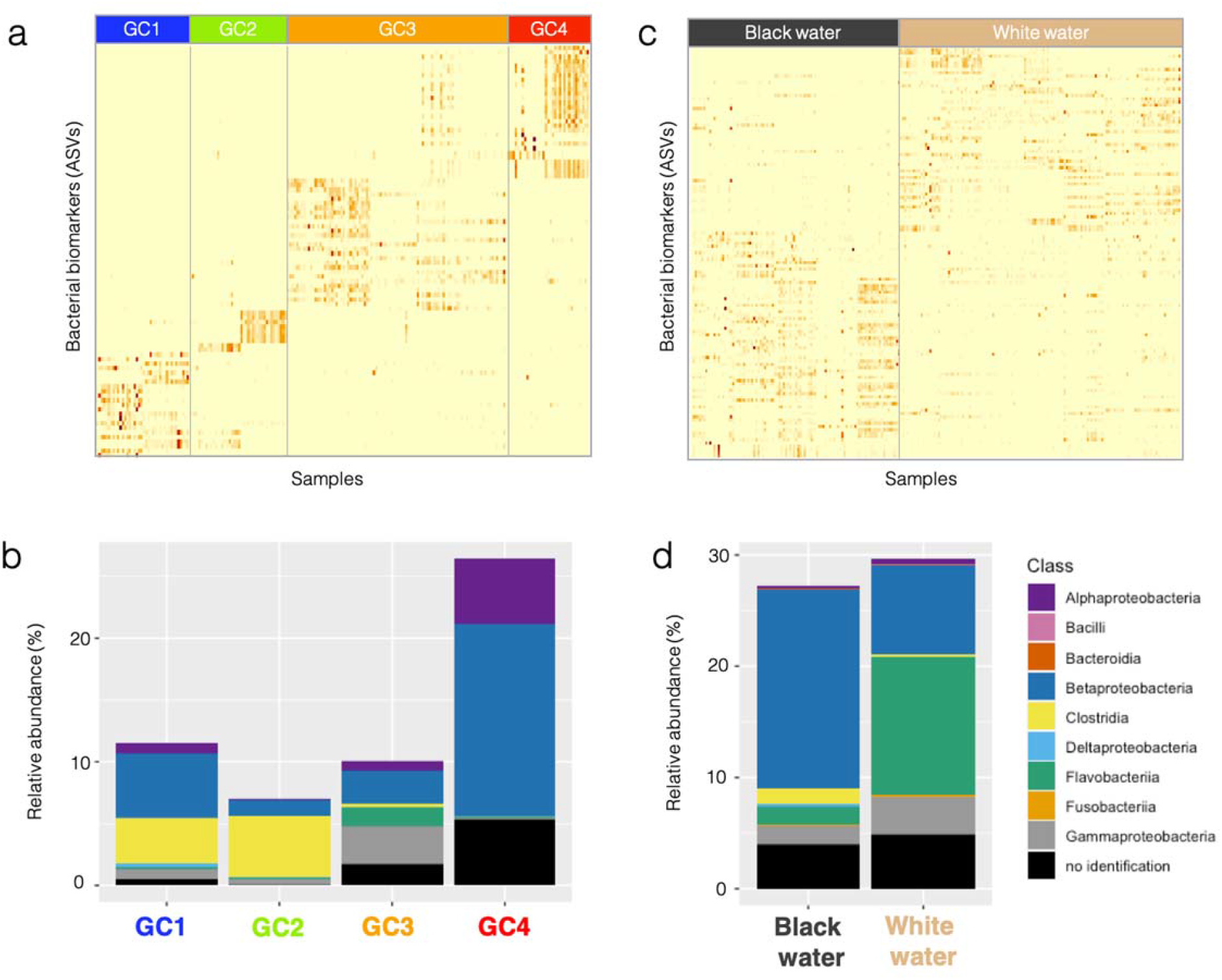
Bacterial biomarkers (at the ASV level) specific to one of the four genetic clusters (a, b) or one of the water types (c, d). In (a, c) the relative abundance of these biomarkers in each genetic cluster (a) or water type (c) is shown on heatmaps where each column is a gill microbiome sample and each row represents one of the biomarker ASV. Darker shades of orange indicate higher relative abundance of the ASV in the sample. In (b, d) the relative abundance of the biomarkers in each genetic cluster (b) or water type (d) is shown on stacked barplots. In these plots, biomarker ASVs are colored according to their phylogeny (Class level).

Concerning the bacterial biomarkers specific to water types, a total of 125 ASVs were identified: 69 ASVs associated to black water and 56 ASVs to white water flag cichlids. Biomarkers specific to black water fish were mostly from the Betaproteobacteria, Deltaproteobacteria and Clostridia classes, while those specific to white water fish were characterized by a relatively higher abundance of Flavobacteriia and Gammaproteobacteria. The biomarkers specific to water types represented an average of 28.4% of the global gill microbiome (27.2% for black and 29.6% for white water fish).

### 2.2 Gill and bacterioplankton communities in an environmental gradient

After sequencing, merging and quality filtering, 1,137,185 reads were obtained (mean of 16,973 reads/sample, N = 67 samples) for the bacterioplankton dataset. Bacterioplankton samples showed a high relative abundance of Proteobacteria (mean of 57 ± 4 %), Actinobacteria (24 ± 3 %) and Firmicutes (17 ± 3 %) transcripts (Fig. 5a). Bacterioplankton samples from black water environments were significantly enriched in Proteobacteria (68 ± 7 % in black water, 49 ± 4 % in white water, Student T test pvalue = 0.02), while bacterioplankton samples from white water environments were significantly enriched in Actinobacteria (29 ± 3 % in white water, 18 ± 4 % in black water, Student T test pvalue = 0.04). A detailed characterization of these bacterioplankton communities can be found in Sylvain *et al*. (2021).

**Figure 5:**
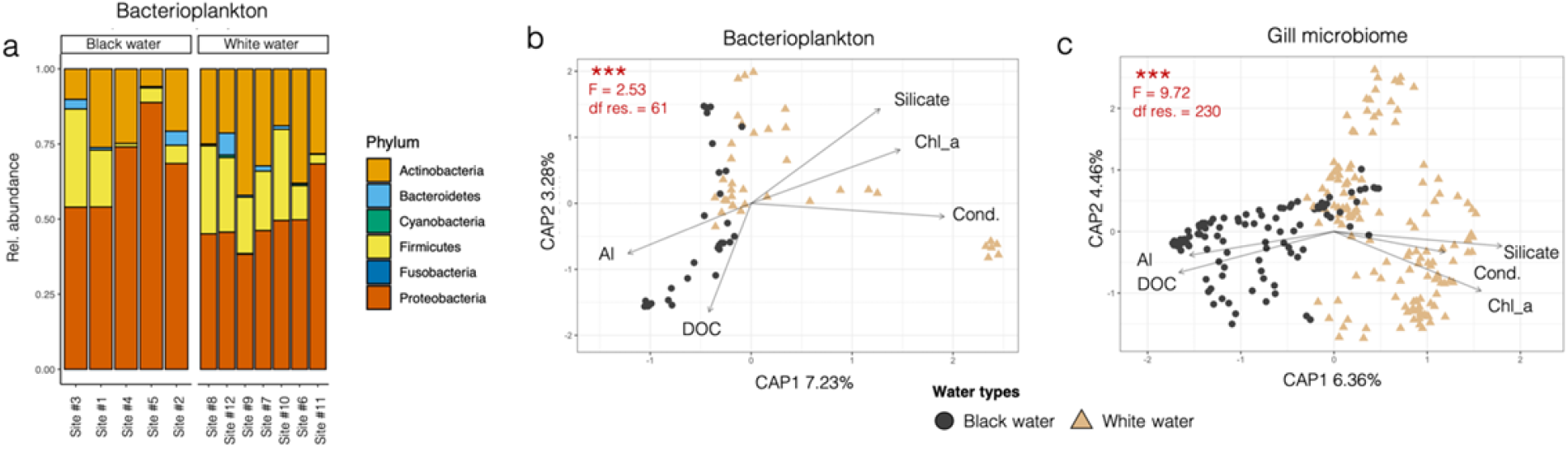
(a) Relative abundance of the main phyla detected in bacterioplankton samples. In (b, c): Constrained analysis of principal coordinates (CAP) on bacterioplanktonic communities (b) and on gill microbiome samples (c). Each data point in the CAP plots represents a sample, and their color and shape correspond to the water type at the sampling site. The results of permutation tests for CAP under reduced model (five environmental variables) are shown in red in the upper left corner of each plot. “df res” stands for residual degrees of freedom and “***” for “pvalue < 0.001”. “Al” stands for dissolved aluminum (μg L-1), “DOC” for the concentration of dissolved organic carbon (mg L-1), “Silicate” for the concentration of silicates (mg L-1), “Chl_a” for the concentration of chlorophyll a (μg L-1), “Cond.” for the conductivity (μS)._

The environmental parameters measured in this study and used to compute CAP constrained ordinations (Fig. 5b,c) indicate that the 12 sampling sites have two contrasting physico-chemical profiles that are typical of black and white water types (Fig. 1b, Suppl. Tables 2, 3, 4, 5, and Suppl. Fig. 5). Additional physico-chemical characterization of these sampling sites can be found in Suppl. Mat. section “Environmental characterization” and in Sylvain *et al*. (2021).

The CAP ordinations (Fig. 5b, c) revealed that samples cluster according to their water type of origin for gill and bacterioplankton communities. Permutation tests for CAP under reduced model showed that the samples significantly clustered according to the five environmental variables provided for the constrained ordination, both for bacterioplankton (pval < 0.001, F = 2.53, df.res = 61) or gill microbiome (pval < 0.001, F = 9.72, df.res = 230). The percentages of the variance retained by the first two dimensions of the CAP were relatively low: 10.51% for bacterioplankton and 10.82% for gill microbiome. This significant difference between communities from different water types was confirmed by PERMANOVA results (bacterioplankton: pval < 0.001, F = 3.51, df.res = 65; gill microbiome: pval < 0.001, F = 14.59, df.res = 234). Fitting of selected environmental variables on CAP (Fig. 5b,c) and *envfit* results (Suppl. Table 6) indicated that variations in the concentrations of Al and DOC were significantly correlated to gill microbiomes from black water sites (Al: R^2^ = 0.17, pval < 0.001; DOC: R^2^ = 0.09, pval < 0.001), while conductivity, concentration of chlorophyll a and silicates were associated with gill microbiomes from white water sites (cond.: R^2^ = 0.17, pval < 0.001; chl. A: R^2^ = 0.23, pval < 0.001; silicate: R^2^ = 0.11, pval < 0.001). Interestingly, the bacterioplankton was not sensitive to the same physico-chemical parameters, as the *envfit* analysis points out: Instead of being correlated with the five parameters above, planktonic communities were significantly correlated with several parameters associated with DOC characteristics, as well as the concentration of phaeopigments, Cr, Fe and Cu (scores in Suppl. Table 6). Gill microbiome samples displayed a more pronounced difference between water types than bacterioplankton samples. However, this could be due to the uneven number of samples (N gills = 236, N bacterioplankton = 67).

### 2.3 Modelling environmental and genetic effects on gill microbiome beta-diversity

We used a LMER modelling approach to identify the main factors, or combination of factors, which explain the bêta-diversity patterns (BC distances) observed on flag cichlid gill microbiomes with the most precision and parsimony (Fig. 6). Spearman correlations between the factors considered in the LMER models varied between 0.004 and 0.31, the highest being between geographical distances and pairwise fixation indexes (F_ST_) between fish from different sampling sites (Suppl. Fig. 4). The dissimilarity values predicted by the global model (Fig. 6a), including all explicative variables, were significantly correlated to the observed data (BC distances) from the fish gill microbiomes (R^2^ = 39.8%, F = 42.3, pval < 0.001). The eight models with Akaike weights > 0, including the null model, are shown in the diagram on Fig. 6b. Of these, only three significant models emerged, and they included only two out of the four explicative variables included in the global model: BC distance among bacterioplankton samples and F_ST_ values between fish from different sites. The models which included the geographical distance between sampling sites and the environmental parameters (Euclidean distances) were not significant as they displayed Akaike weights inferior to the null model. The model with the best score was only composed of the bacterioplankton BC distance variable (Akaike weight = 0.51, K = 5, AICc = −161.15), largely surpassing the second-best model, which only included the F_ST_ values (Akaike weight = 0.14, K = 5, AICc = −158.55). The models including the bacterioplankton BC distance variable (Fig. c) had a model-averaged estimate of 0.45 (conf. interval 0.12 - 0.79), more than three times the estimate for models including the F_ST_ values (0.13 with conf. interval 0.02 – 0.24) (Fig. d). When linear correlations were plotted separately, outside of the LMER model (Fig. c, d, e, f) they showed that out of the four explicative variables considered, the bacterioplankton BC distance variable was the most strongly correlated to gill microbiome dissimilarity (Fig. 5c; R^2^ = 9.5%, F = 6.7, pvalue < 0.05), while Euclidean distances between environmental parameters were the least correlated to gill microbiome dissimilarity (Fig. 5f; R2 = 1.0%, F = 0.6, pvalue = 0.4). Overall, the mixed-effects linear modelling analysis suggested that bacterioplankton dissimilarity was much more correlated to gill microbiome dissimilarity than the host F_ST_ values, and the two other explicative variables considered.

**Figure 6:**
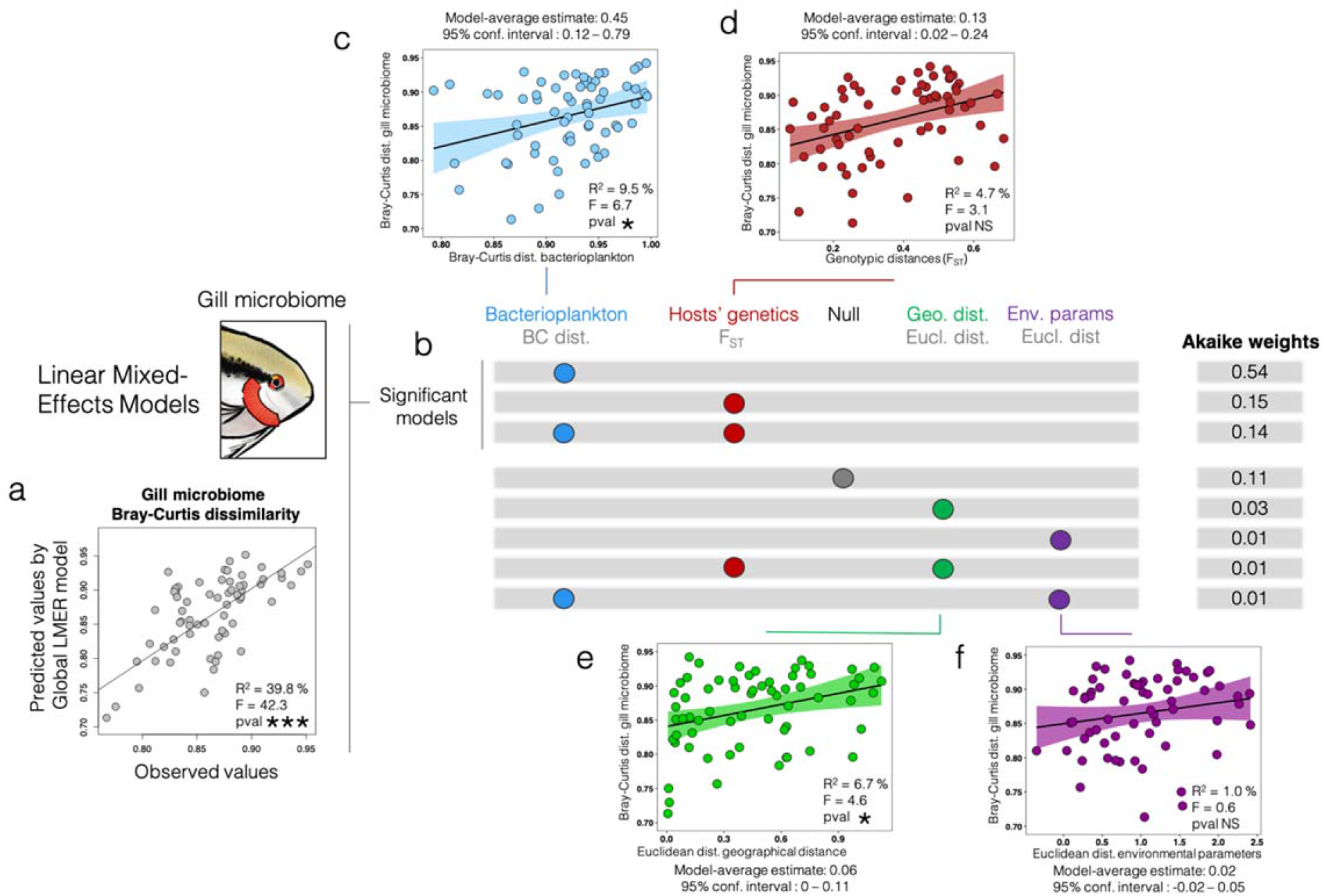
Linear mixed-effects modeling analysis of the explicative variables potentially associated to gill microbiome transcriptional activity. The four following explicative variables (transformed in distance matrices) are included in the global model (a) used for predicting dissimilarity values: bacterioplankton transcriptional activity (BC distance), genotypic distances between hosts (F_ST_), geographic distances between sites (Euclidean distance), and the ensemble of the 34 environmental parameters measured in this study (Euclidean distance). Each row of the gray diagram in (b) corresponds to one of the models tested, and colored circles represent the elements that were included in the models. The upper three rows of the diagram represent significant models, with Akaike scores superior to the null (random) model (fourth row), while the lower four rows correspond to models with low explicative power. Linear correlation plots (c, d, e, f) for each of the four explicative variables considered in the model represent the linear correlation between the normalized distance/dissimilarity of each variable with the gill microbiome Bray-Curtis (BC) dissimilarity, when assessed separately outside of the LMER model. The linear correlations with BC gill microbiome dissimilarity are displayed with the following variables: (c) BC bacterioplankton dissimilarity, (d) F_ST_ genotypic distances, (e) Euclidean distances of geographical distances, (f) Euclidean distances of environmental parameters.

## 3. Discussion

Fish-microbe systems evolve according to a complex mixture of environmental and host-specific factors (Llewellyn *et al*. 2014). External microbiomes of Amazonian fish are known to be especially sensitive to environmental factors, most notably to the composition of bacterioplankton (Sylvain *et al*. 2020). Several studies have investigated the Amazonian bacterioplankton composition, from the upper (Toyama *et al*. 2016, Santos-Junior *et al*. 2017; Sylvain *et al*. 2019, 2021) to the lower Amazon and its plume in the Atlantic Ocean (Satinsky *et al*. 2014ab, 2015; Doherty *et al*. 2017). The bacterioplankton communities characterized in our study were in accordance with results from previous studies: Proteobacteria, Actinobacteria, Firmicutes and Bacteroidetes were also the main phyla identified by Toyama *et al*. (2016) and Santos-Junior *et al*. (2017). In addition, these two investigations reported the occurrence of Cyanobacteria (up to 24% in certain samples) and Planctomycetes (up to 8%) which were also detected but were not as abundant in our study. These differences could be due to the fact that sampling sites visited in these investigations were different from ours, and that our molecular approach was focused on the detection of transcriptionally-active taxa only.

To our knowledge, the present study is the first to characterize the gill microbiome of an Amazonian fish. Several taxonomic groups detected on the gills were also found in the skin mucus of flag cichlids in a previous investigation (Sylvain *et al*. 2019). In both communities, Proteobacteria, Actinobacteria and Bacteroidetes were among the most abundant phyla. Our results are also in accordance with a previous investigation describing the gill microbiome from 53 reef fish species, which identified Proteobacteria, Firmicutes, Fusobacteria and Bacteroidetes as one of the most abundant phyla on fish gills (Pratte *et al*. 2018). Furthermore, our results (Fig. 3b, Fig. 5c) show a strong response of the flag cichlid gill microbiome to the water type. This response could depend on the different physico-chemical conditions, characteristic of each water type, but also on the host physiological response to these conditions. For instance, a study on the gills of the Amazonian sardine (*Triportheus albus*) has shown a contrasted regulation of physiological processes in different water types (Araújo *et al*. 2017). Some of these processes, such as the regulation of tight junction permeability, metal ion binding and the host immune response (Araújo *et al*. 2017), potentially involve gill microbial communities (Ulluwishewa *et al*. 2011; Becker and Skaar 2014; Stevens *et al*. 2021).

The analysis of gill bacterial biomarkers has proven to be an essential tool in differentiating fish from different environmental or health conditions (Legrand *et al*. 2017). Here, the biomarkers’ analysis (Fig. 4) highlighted the potential of several bacterial groups from the gills to distinguish fish from different water types. Betaproteobacteria were mostly associated with black water fish (Fig. 4d). Interestingly, a recent study on Atlantic salmon showed a major enrichment of Betaproteobacteria in microbiomes characterized from whole gill samples instead of gill swabs (Clinton *et al*. 2021). This former study suggests that the localization of selected members of the Betaproteobacterial group could be within cryptic tissue locations, such as beneath the surface epithelium. A few strains of Betaproteobacteria are known to be endosymbionts of fish gills (Mitchell *et al*. 2013; Gjessing *et al*. 2021). Accordingly, in white water environments we detected several ASV biomarkers that were from the Flavobacteriia class, mostly represented by members of the genus *Chryseobacterium*, another clade known to include potential intracellular bacteria (Lim *et al*. 2020). In the future, using a correlative imaging approach, as done on mussel gills by Franke *et al*. (2021), would help to resolve the potential niche partitioning of variable microbial communities across the ultrastructure of gill tissue and benefit our understanding of the interaction between these water-type biomarkers and their fish host in different habitats.

### Decomposing the factors driving the gill microbiome

Not only the gills’ microbiome (Fig. 5c), but also the bacterioplankton activity (Fig. 5b) and the environmental physico-chemical parameters (Suppl. Fig. 5) varied significantly varied according to water type. Gill microbial communities are located at the frontier between the host and the external environment. This unique location enables them to interact extensively with their host and the metabolites of the bacterioplankton community, but it also exposes them to the variations of water physico-chemical parameters. For this reason, we decomposed the effect of the water type into two factors in the LMER model: the environmental physico-chemical parameters, and the bacterioplankton.

#### Environmental physico-chemical parameters

It is well known that water physico-chemical parameters drive, to a certain extent, the composition and expression of aquatic microbial communities (Pang *et al*. 2014; Lindh and Pinhassi 2018; Fadeev *et al*. 2021). This has also been shown previously in a study focused on Amazonian bacterioplankton (Sylvain *et al*. 2021). In our investigation, when all environmental physico-chemical parameters were considered together to build distance matrices, the output was not significantly associated to patterns observed on the gill microbiomes (Fig. 5). However, when considered separately in the CAP constrained ordinations, they did show significant correlation to gill microbiome beta-diversity (Fig. 5c). Results from the *envfit* analyses (Suppl. Table 6) also suggested a significant correlation between selected physico-chemical parameters and the dissimilarity patterns observed for the gill microbiomes. The results thus suggested that flag cichlid gill microbiome communities were sensitive to few (but not all) of the parameters associated with water type, most notably conductivity, Al concentrations, silicate, DOC and chlorophyll a (Fig. 5c). Accordingly, previous studies have observed a significant influence of conductivity (Krotman *et al*. 2020), chlorophyll a (Minich *et al*. 2020) and DOC concentrations (Sylvain *et al*. 2020) on fish skin mucus bacterial communities.

#### Bacterioplankton

Among the factors considered in the modeling analysis, the bacterioplankton dissimilarity was the best explicative variable that correlated to the gill microbiome dissimilarity (Fig. 6b, c). This result makes sense physiologically, as in this case, both the explicative and the response variables were bacterial communities. Several studies have shown that microbial communities from different microhabitats (e.g. from different hosts sharing the same environment), can show similar responses when exposed to the same environmental conditions. For instance, studies on corals have shown that stressors such as lowered pH or eutrophication often lead to an increase in coral microbiome alpha diversity and a decrease in the bacterial symbiont *Endozoicomonas*, a pattern consistent across different coral species and geographical regions (reviewed in McDevitt-Irwin *et al*. 2017). In our study system, it is possible that direct interactions exist between both bacterial community types considered here: Bacterioplankton transcriptional activity could affect the activity of bacterial members from the gill microbiome. For example, recent investigations suggested that the production of bioactive compounds, such as cyanotoxins produced by planktonic cyanobacterial strains, can significantly perturb fish microbiomes (Gallet *et al*. 2021; Duperron *et al*. 2019).

An additional process linking planktonic and fish-associated microbiomes relies in the assembly process of the later. Indeed, the assembly of fish microbiomes largely depends on the past and present environmental pools of bacterial taxa (Llewellyn *et al*. 2016; Sylvain *et al*. 2017, Lavoie *et al*. 2021). For instance, a study on Atlantic salmon has showed that the external skin mucus microbiome changes when the environment is modified: The microbiome taxonomic structure of hatchery-raised juveniles converged with their wild counterparts shortly after transferring them from the hatchery to the wild (Lavoie *et al*. 2021). The importance of “source-sink” processes has also been shown in the microbial communities’ assembly of the Amazonian discus fish, where bacterioplanktonic taxa shape the gut microbiome at early life stages, a period during which larvae microbiomes show a weak resistance to colonization (Sylvain and Derome 2017). The importance of source-sink processes might be even more important for gill communities, as they are exposed to environmental stressors, just like fish skin mucus communities. Such external communities are known to have an especially dynamic composition that varies in accordance to fine-scale changes in environmental conditions (Sylvain *et al*. 2016; Audet-Gilbert *et al*. 2021) and are thus more susceptible to frequent remodeling by environmentally-recruited taxa. Finally, while direct interactions between bacterioplankton and gill microbiome could be the cause of the correlation between these communities in our datasets, there is also a possibility that both communities show similar responses to shared water physico-chemical parameters without interacting directly with one another. However, based on the close physical proximity of these communities and the high recruitment potential of environmental strains by fish microbiomes, this perspective seems unlikely.

Overall, the high sensitivity of gill communities to the water type could potentially be beneficial for fish holobionts in Amazonian ecosystems. Indeed, the rapid remodeling of gill communities could be an asset in heterogenous systems such as the Amazon basin, as it could facilitate the adaptation of fish to changing conditions, for instance during the important seasonal floods occurring in the flag cichlid’s habitat during the wet season (January to July). Furthermore, since microbial communities are known to play important roles in membrane permeability (reviewed in Ghosh *et al*. 2021), they are likely involved in the regulation of ionoregulatory processes occurring at the gills. In that case, a dynamic gill microbiome that adapts rapidly to changing environmental conditions could be an essential tool to optimize ionic balance, especially in mixed-water environments receiving inputs of both ion-rich white waters and ion-poor black waters. Metatranscriptomic results from other Amazonian fish species would be needed in the future to test this hypothesis.

#### The host’s genetic effects

The second-best model to explain the gill microbiome bêta-diversity patterns was the one including only the pairwise fixation indexes (F_ST_) (Fig. 6b). Thus, a significant influence of the host’s genetics on the gill microbiome is not to be completely excluded, even though the best model only included bacterioplankton dissimilarity values. Indeed, the selection of the bacterioplankton model based on Akaike scores gave a lot of weight to the parsimony criteria, thus advantaging simpler models (Bates *et al*. 2015). Realistically, the global model was the one making the most sense biologically, as it included the influence of all the ecological, physico-chemical and geographical explicative variables considered in this study, which are known to affect the dynamics of host-associated microbiomes in many ecosystems (Goertz *et al*. 2019; Sylvain *et al*. 2020, 2021; Gallet *et al*. 2021). It may seem surprising that the F_ST_ values did not show a significant linear correlation with gill microbiome beta-diversity (Fig. 6d), when this explicative variable alone constituted the second-best model (Fig. 6b). This result is likely due to the fact that the linear correlation on Fig. 6d did not account for the influence of the other explicative variables which potentially also influenced gill microbiome communities, while LMER models do. The LMER models thus enabled us to partition the effect of various variables acting simultaneously on gill communities.

Our study was one of the first investigations that combined microbiome data to host’s genetic information collected on the same fish specimens. Overall, gill microbiome samples were significantly different according to their genetic cluster of origin (Fig. 3a), suggesting that the flag cichlid genetic background might influence its microbiome transcriptional activity. However, three main results suggest that the effect of the host genetics on the transcriptional activity of gill microbiomes was weak in comparison to environmental factors. Firstly, Fig 3b highlights that for the two genetic clusters that contained sites of both water types (GC2 and GC4), the differentiation between gill microbiome samples of different water types was stronger than between samples from different genetic clusters. The PERMANOVA tests on this figure resulted in higher F and R^2^ values in Fig. 3b than in Fig. 3a, even though the number of residual degrees of freedom was lower for both genetic clusters in Fig. 3b. The percentage of variance explained by the first two axes of PCoAs in Fig. 3b were also higher than those from Fig. 3a. Secondly, the analysis of bacterial biomarkers (Fig.4) revealed that biomarkers associated to water type constituted a higher relative proportion (125 ASVs representing an average of 28.4%) of the total microbiome, than biomarkers associated to the genetic clusters (90 ASVs representing an average of 13.8% of the total microbiome). Thirdly, the relatively low Akaike score of the best LMER model including the host pairwise fixation index (i.e. three times inferior to the best one including bacterioplankton BC distance), as well as the relatively low model-average estimate of this factor (0.13) also suggest a weak correlation between host genetics and gill microbiome transcriptional activity. A weak influence of host genetics on gill microbiomes has also been reported in different genetic lines of rainbow trout (Brown *et al*. 2019). Additionally, a recent study focused on the external skin mucus microbiome of teleosts and elasmobranchs found that associations between hosts genetics and their microbiome composition was clade-specific (Doane *et al*. 2020), thus the patterns observed on the flag cichlid microbiome might differ in other Amazonian fish.

## 4. Conclusions

This study on the flag cichlid gill microbiome is one of the first to include information on the hosts’ genetic backgrounds, the free-living bacterioplanktonic pools of bacteria and environmental parameters. These datasets enabled us to study the relative contribution of genomic and environmental factors shaping the flag cichlid gill microbiome transcriptional activity in a simple natural system including fish from four genetic clusters and two contrasting Amazonian water types. We observed that gill microbiome samples were significantly different between the genetic clusters and between water types, suggesting a significant contribution of both host- and environment-specific factors in shaping these bacterial communities. However, constrained ordinations, PERMANOVAs and analyses of bacterial biomarkers suggest that the contribution of the host’s genetic background was relatively weak in comparison to environment-related factors in structuring the gill communities. This result was also confirmed by a mixed-effects modeling analysis, which suggested that the dissimilarity among bacterioplanktonic communities possessed the highest explicative power regarding the dissimilarity of gill microbiomes, while pairwise fixation indexes (F_ST_) from hosts only had a weak explicative power. Whether the relative importance of the host’s genomic background on the gill microbiome is clade-specific should be confirmed in future studies targeting a broader range Amazonian species and, if possible, using a reciprocal transplant experiment in controlled environments.

## 5. Materials and Methods

### 5.1 Ethical approval

This study was carried out in accordance with the recommendations of the Ethics Committee for the Use of Animals of the *Instituto Nacional de Pesquisas da Amazonia* (INPA). The permit (number 29837-18) was approved by the Ethics Committee for the Use of Animals of INPA.

### 5.2 Fish sampling

A total of 240 fish were collected from 12 sampling sites distributed throughout the upper Brazilian Amazon Basin in October-November 2018 and 2019 (dry season). Sampling fish during the dry season significantly facilitated the fish collection and enabled us to reach our objective of 20 specimens per sampling site. GPS coordinates and a map of all sites are found in Suppl. Table 1 and Fig. 1a, respectively. Over the 12 sampling sites, there were seven white water sites and five black water sites. Twenty fish specimens were collected at each sampling site using a combination of small seine net-fishing and line fishing. One gill arch was sampled from each specimen immediately after collection using sterile dissection tools (EtOH 70%). The biological samples were preserved in 2 mL of NAP conservation buffer to preserve RNA integrity (Camacho-Sanchez *et al*. 2013; Menke *et al*. 2017) and were then stored at −80°C until processing.

A fin clip was also sampled on the same specimens for the phylogenomic investigation of Leroux *et al*. (2022). The genetic cluster (GC) of specimens sampled at each sampling site, previously determined by Leroux *et al*. (2022), is detailed in Fig. 1, Suppl. Fig. 1 and Suppl. Table 1.

### 5.3 Bacterioplankton sampling

Six water samples were collected per site to characterize the transcriptional activity of the bacterioplankton community. Surface water samples were taken at a depth of 30 cm in 2 L Nalgene^TM^ (Thermo Fisher Scientific, Waltham (MA) USA) bottles. Filtration was performed as in Cruaud *et al*. (2017) through 22 µm-pore size polyethersulfone Sterivex^TM^ filters (cat #SVGP01050, Millipore, Burlington (MA), USA) less than 30 minutes after collection. Filters were also stored in 2 mL of NAP conservation buffer immediately upon collection, and then stored at −80°C until processing. Before RNA extraction, Sterivex^TM^ filter casings were opened and processed according to Cruaud *et al*. (2017) using sterile instruments, and filter membranes were stored in TRIzol^TM^ (cat #15596026, Thermo Fisher Scientific).

### 5.4 Characterization of environmental variables

A total of 34 environmental variables commonly characterized in limnological studies (Cohen *et al*. 2020) and associated to the physicochemical differences between the two water types (Sioli 1984) were measured (Suppl. Tables 2, 3, 4, 5). The methods used to measure the environmental variables are detailed in Sylvain *et al*. 2021. In brief, temperature (°C), conductivity (µS), pH and dissolved oxygen (%) were measured directly on site using a YSI professional plus series multimeter (YSI Inc/Xylem Inc, Yellow Springs (OH), USA). At the laboratory, we measured the concentration of DOC, dissolved metals, nutrients, free ions and chlorophyll a also as described in Sylvain *et al*. (2021). The water type (Suppl. Table 1) of each site was determined based on the physicochemical profiles of the sampled environments (Fig. 1b, Suppl. Tables 2, 3, 4, 5). Our measurements of environmental parameters confirmed the *a priori* knowledge of the water types of these sampling sites.

### 5.5 Biological sample processing

#### Gills and bacterioplankton

RNA extractions of whole gills and Sterivex^TM^ filters (bacterioplankton) were performed according to the manufacturer’s instructions of TRIzol^TM^ without modification. Using RNA extracts prevented the detection of inactive, dead or dormant taxa which are normally detected with a standard 16S approach based on DNA extracts. Four blank controls of NAP buffer were also processed identically to all samples for RNA extractions and sequencing. Microbial community transcriptional activity was assessed using a 16S rRNA approach conducted on RNA extracts. The RNA retrotranscription was performed with the qScript cDNA synthesis kit (cat #95048-100) from QuantaBio (Beverly (MA), USA) according to the manufacturer’s instructions. Then, the fragment V3-V4 (∼500 bp) of the 16S rRNA gene was amplified in cDNA extracts by PCR using the forward primer 347F (5’-GGAGGCAGCAGTRRGGAAT-3’) and the reverse primer 803R (5’-CTACCRGGGTATCTAATCC-3’) (Nossa *et al*. 2010). The first PCRs of gill DNA were conducted in 25 μl according to the manufacturer’s instructions of Q5^®^ High-Fidelity DNA Polymerase from New England BioLabs^®^ Inc (cat # M0491S) using an annealing temperature of 64°C and 35 amplification cycles. A PCR protocol with higher sensitivity was used for bacterioplankton samples due to their lower average cDNA concentration: PCRs of bacterioplankton cDNA were performed in 25 μl according to the manufacturer’s instructions of the QIAGEN® Multiplex PCR kit (cat #206143) using an annealing temperature of 60°C and 30 amplification cycles. A second PCR was done using Q5^®^ High-Fidelity DNA Polymerase for both gill and bacterioplankton libraries (12 cycles), to add barcodes (indexes); more details in Suppl. Mat. section “Details on 16S rRNA library preparation”. After each PCR, the amplified DNA of gills and bacterioplankton was purified with AMPure beads (cat #A63880, Beckman Coulter, Pasadena (CA), USA), according to the manufacturer’s instructions, to eliminate primers, proteins, dimers, and phenols. Post-PCR cDNA concentrations were assessed on a Qubit^TM^ instrument (Thermo Fisher Scientific) and by electrophoresis on 2% agarose gels. After purification, multiplex sequencing was performed on Illumina MiSeq by the *Plateforme d’Analyses Génomiques* at the *Institut de Biologie Intégrative et des Systèmes* of Université Laval.

The R package *DADA2* (Callahan *et al*. 2016) was used for amplicon sequence variant (ASV) picking. Quality control of reads was done with the *filterAndTrim* function using the following parameters: 290 for the forward read truncation length, 270 for the reverse read truncation length, 2 as the phred score threshold for total read removal, and a maximum expected error of 2 for forward reads and 3 for reverse reads. The filtered reads were then fed to the error rate learning, dereplication, merging and ASV inference steps using the functions *learnErrors*, *derepFastq*, *mergePairs* and *DADA* from the *DADA2* pipeline (Callahan *et al*. 2016). Chimeric sequences were removed using the *removeBimeraDenovo* function with the *consensus* method parameter. Sequenced PCR negative controls were used to remove ASVs identified as potential cross contaminants using the *isContaminant* function from the *decontam* package with the default threshold of 0.4. Taxonomic annotation of amplicon sequence variants (ASV) was performed by using *blastn* matches against NCBI “16S Microbial” database. As the NCBI database for 16S sequences is larger and updated more frequently than other sources, it provides more information about lesser-known taxa while minimizing ambiguous annotations. Matches above 99% identity were assigned the reported taxonomic identity. Sequences with no match above the identity threshold were annotated using a lowest common ancestor method generated on the top 50 blastn matches obtained, a method inspired from the LCA algorithm implemented in MEGAN (Huson *et al*. 2016). Analyses of Shannon diversity according to sampling depth for each sample are provided in Suppl. Fig. 1 and 2. ASV tables, metadata files and taxonomy information were incorporated into *phyloseq*-type objects (McMurdie and Holmes 2013) before downstream analyses.

#### Fin samples

Detailed methods on the molecular approach, the construction of Genotyping-By-Sequencing libraries, sequencing and sequences’ processing are found in Leroux *et al*. (2022).

### 5.6 Data availability

The 16s rRNA datasets generated and analysed during the current study can be found in the Sequence Read Archive (SRA) repository, BioProjectIDs: PRJNA839167 and PRNJA839174 (data from gills), PRJNA736442 and PRJNA736450 (data from bacterioplankton). The R scripts used for the 16S RNA sequence analysis and the input files including all metadata and ASV taxonomy are freely available on the Open Science Network platform (URL: https://osf.io/d3n76/).

### 5.7 Statistical analyses

The statistical analyses describing the four genetic clusters of flag cichlids used in this study are detailed in Leroux *et al*. (2022) and their results are summarized in Suppl. Fig 1. Using stacked barplots (Fig. 2), we first visualized gill microbiome transcriptional activity according to the water type and the genetic cluster of the host fish. We then used principal coordinates analyses (PCoA) to study how the samples cluster according to the two aforementioned factors (Fig. 3). To compute PCoAs, we used a matrix of pairwise Bray-Curtis (BC) distances between all the gill microbiome samples (N = 240). The same BC distance matrix was used to compute permutational analyses of variance (PERMANOVA) to assess if gill microbiome samples significantly differed according to their water type or the genetic cluster of the host fish (Fig. 3). PERMANOVAs were computed with the *adonis* function of the from *vegan* R package (Oksanen *et al*. 2020) using 1000 permutations.

Secondly, we identified bacterial biomarkers (at the ASV level) from the gill microbiome, that were significant discriminant features of host genetic cluster (Fig. 4a, b) or water type (Fig. 4b, c). To do so, we used the function *multipatt* from the *IndicSpecies* package (De Caceres and Legendre 2009) to identify discriminant ASVs based on their read abundance in the gill microbiome ASV table. The ASVs which showed indicator values > 0.5 and p-values < 0.01 after 999 permutations were retained as discriminant features for one of the groups from the comparison (i.e. the four genetic clusters or the two water types). The indicator value used for biomarker identification incorporates a correction for unequal group sizes (Dufrêne and Legendre 1997). The relative abundance of these biomarkers was represented in heatmaps (Fig. 4a, c) and stacked barplots (Fig. 4b, d).

Thirdly, we studied in more detail how the transcriptional activity of gill bacterial communities varied according to two environment-related factors: Bacterioplankton transcriptional activity and environmental parameters (Fig. 5). To do so, we first represented the relative abundances of transcripts from the different bacterial phyla detected in bacterioplankton communities using stacked barplots (Fig. 5a). Then, we used the *capscale* function from vegan R package (Oksanen *et al*. 2020) to compute constrained analyses of principal coordinates (CAP), a distance-based ordination method similar to redundancy analyses (Legendre and Anderson 1999), to summarize the variation in the bacterioplankton or gill microbiome (response variables) explained by environmental parameters (explanatory variables) (Fig. 5b,c). BC distance was used for the computation of CAPs. Covariation and multicollinearity were assessed by measuring variance inflation factors (VIF) (Blanchet *et al*. 2008) using the function *vif.cca*. Only environmental parameters not significantly correlated to each other, with VIF scores < 10 were kept as explanatory variables. Consequently, the five following parameters were retained for this analysis: water conductivity (µS/L), and the concentrations of DOC (mg/L), aluminum (µg/L), silicate (µmol/L) and chlorophyll a (µg/L) (Fig. 5b, c). Permutation tests (999 permutations) for CAP under reduced model (five environmental variables) were conducted by running *anova* on the *capscale* result. The function *envfit* from *vegan* (Oksanen *et al*. 2020) was used to measure the strength of association between environmental variables and microbiome data (from the relative abundance table of ASVs). PERMANOVAs based on BC distances were computed to detect significant differences between samples of different water types.

Finally, we built linear mixed-effects (LMER) models from ecological, physio-chemical and geographical variables, that could potentially affect the gill microbiomes’ transcriptional activity measured in this study (Fig. 6). These factors included: the bacterioplankton transcriptional activity, host fish genetics, geographical distance between sampling sites, and the environmental parameters measured. Here, the factor “Hosts’ genetics” does not refer to the four genetic clusters considered in Fig. 2, 3, 4, but rather to the list of single nucleotide polymorphisms (SNP) used to compute pairwise fixation indexes (F_ST_) with the LMER model. Details on the computation of F_ST_ values are found in Leroux *et al*. (2022). The covariation between these four factors was assessed using Spearman correlations, before model construction (Suppl. Fig. 4). Our LMER model was based on distance matrices, thus the following distances were calculated between samples of different sampling sites: (1) BC distance for bacterioplankton and gill microbiome transcriptional activity; (2) F_ST_ were estimated for host fish genetic dissimilarity; (3) Euclidean distances were computed (after normalizations based on the mean and standard deviation) for geographical distances, and for the set of environmental parameters. We identified the combination of factors which most likely and parsimoniously explain the patterns observed in the bêta-diversity (BC distance) of gill microbiome transcripts (Douglas *et al*. 2015), using the model selection tool *aictab* from the R package *AICcmodavg* (Buckland *et al*. 1997; Burnham and Anderson 2002), based on Akaike weights. A global model containing all potential factors (Fig. 6a) and a null model (random vector) were also included in the analysis. The fitting of the models to gill microbiome data was done with *lmer* from *lme4* on R (Douglas *et al*. 2015). Model-averaged parameter estimates, the unconditional standard error and unconditional confidence intervals were computed using *AICcmodavg* (Buckland *et al*. 1997; Burnham and Anderson 2002). Linear correlation of individual variables and the Bray-Curtis distances of the gill microbiomes were also plotted separately (Fig. 6c, d, e,f), outside of the LMER models.

## Acknowledgements

We thank the National Geographic Society, *NSERC*, MITACS, and Ressources Aquatiques Québec for awarding travel and field work grants to FÉS. This study was part of the NSERC Discovery grant of ND, the INCT ADAPTA project of ALV, and supported by a Canada-Brazil Awards – Joint Research Project of ND and ALV, by CNPq, FAPEAM and CAPES. We thank Thiago Nascimento, Reginaldo Oliveira and Nazaré Paula for technical support with field work logisitcs. We thank Roxanne Dhommée for support in the molecular biology laboratory work. Finally, we thank the anonymous reviewers that generously took the time to help improve this manuscript. No conflicts of interests.

## Authors’ contributions

F-ÉS, ALV and ND designed the study; F-ÉS, NL, AH and ND performed field sampling; F-ÉS, NL and PLM conducted RNA extractions and prepared 16S rRNA libraries; F-ÉS, NL and SB conducted the bioinformatical and statistical analyses; F-ÉS wrote the manuscript; all authors revised the manuscript.

## References

Araújo, J., Ghelfi, A., Val, A. L. 2017. *Triportheus albus* Cope, 1872 in the blackwater, clearwater, and whitewater of the Amazon: A case of phenotypic plasticity?. Frontiers in genetics, 8, 114. doi: 10.3389/fgene.2017.00114

Bates, D., Kliegl, R., Vasishth, S., Baayen, R. H. 2015a. Parsimonious mixed models. arXiv, 2015a1506.04967. doi: 10.48550/arXiv.1506.04967

Bates, D., Maechler, M., Bolker, B., Walker, S. 2015. Fitting Linear Mixed-Effects Models Using lme4. Journal of Statistical Software, 67(1), 1–48. doi:10.18637/jss.v067.i01

Becker, K. W., Skaar, E. P. 2014. Metal limitation and toxicity at the interface between host and pathogen. FEMS microbiology reviews, 38(6), 1235–1249. doi: 10.1111/1574-6976.12087

Blanchet, F. G., P. Legendre, D. Borcard. 2008. Forward selection of explanatory variables. Ecology, 89: 2623–2632. doi:10.1890/07-0986.1

Bledsoe, J. W., Waldbieser, G. C., Swanson, K. S., Peterson, B. C., Small, B. C. 2018. Comparison of channel catfish and Bbue catfish gut microbiota assemblages shows minimal effects of host genetics on microbial structure and inferred function. Frontiers in Microbiology, 9. doi:10.3389/fmicb.2018.01073

Bogotá-Gregory, J.D., Lima, F.C.T., Correa, S.B., Silva-Oliveira, C., Jenkins, D. G., Ribeiro, F. R., Lovejoy, N. R. Reis, R. E., Crampton, W. G. R. 2020. Biogeochemical water type influences community composition, species richness, and biomass in megadiverse Amazonian fish assemblages. Scientific Reports, 10:15349. doi: 10.1038/s41598-020-72349-0

Borghezan, E. A., Pires, T. H. S., Ikeda, T., Zuanon, J., Kohshima, S. 2021. A review on fish sensory systems and Amazon water types with implications to biodiversity. Frontiers in Ecology and Evolution, 8:589760. doi : 10.3389/fevo.2020.589760

Boutin, S., Sauvage, C., Bernatchez, L., Audet, C., Derome, N. 2014. Inter individual variations of the fish skin microbiota: host genetics basis of mutualism?. PloS one, 9(7):e102649. doi: 0.1371/journal.pone.0102649

Buckland, S. T., Burnham, K. P., Augustin, N. H. 1997. Model selection: an integral part of inference. Biometrics, 53:603–618.

Brown, R. M., Wiens, G. D., Salinas, I. 2019. Analysis of the gut and gill microbiome of resistant and susceptible lines of rainbow trout (*Oncorhynchus mykiss*). Fish & shellfish immunology, 86:497–506. doi: 10.1016/j.fsi.2018.11.079

Burnham, K. P., Anderson, D. R. 2002. Model selection and multimodel inference: a practical information-theoretic approach. Second edition. Springer: New York.

Callahan, B. J., McMurdie, P. J., Rosen, M. J., Han, A. W., Johnson, A. J. A., Holmes, S. P. 2016. DADA2: High-resolution sample inference from Illumina amplicon data. Nature Methods, 13:581. doi:10.1038/nmeth.3869

Camacho-Sanchez, M., Burraco, P., Gomez-Mestre, I., Leonard, J. A. 2013. Preservation of RNA and DNA from mammal samples under field conditions. Molecular Ecology Resources, 13: 663–673. doi:10.1111/1755-0998.12108

Clinton, M., Wyness, A.J., Martin, S.A.M., Brierley, A. S., Ferrier, D. E. K. 2021. Sampling the fish gill microbiome: a comparison of tissue biopsies and swabs. BMC Microbiology, 21:313. doi: 10.1186/s12866-021-02374-0

Cohen, R. S., Gray, D. K., Vucic, J. M., Murdoch, A. D., Sharma, S. 2021. Environmental variables associated with littoral macroinvertebrate community composition in Arctic lakes. Canadian Journal Fisheries and Aquatic Science, 78:110–123. doi:10.1139/cjfas-2020-0065

Cooke, G. M., Landguth, E. L., Beheregaray, L. B. 2014. Riverscape genetics identifies replicated ecological divergence across an Amazonian ecotone. Evolution, 68: 1947–1960. doi:10.1111/evo.12410

Cruaud, P., Vigneron, A., Fradette, M-S., Charette, S. J., Rodriguez, M. J., Dorea, C. C., Culley, A. I. 2017. Open the Sterivex (TM) casing: An easy and effective way to improve DNA extraction yields. Limnology and Oceanography: Methods, 15:1015–1020. doi:10.1002/lom3.10221

Dal Pont, G., Domingos, F. X. V., Fernandes-de-Castilho, M., Val, A. L. 2017. Potential of the biotic ligand model (BLM) to predict copper toxicity in the white-water of the Solimões-Amazon River. Bulletin of Environmental Contaminants and Toxicology, 98: 27–32. doi:10.1007/s00128-016-1986-1

De Caceres, M., Legendre, P. 2009. Associations between species and groups of sites: indices and statistical inference. Ecology, 90(12):3566–3574. doi: 10.1890/08-1823.1

Doane, M.P., Morris, M.M., Papudeshi, B. et al. 2020. The skin microbiome of elasmobranchs follows phylosymbiosis, but in teleost fishes, the microbiomes converge. Microbiome, 8:93. doi: 10.1186/s40168-020-00840-x

Doherty, M., Yager, P. L., Moran, M. A., Coles, V. J., Fortunato, C. S., Krusche, A. V., Medeiros, P. M., Payet, J. P., Richey, J. E., Satinsky, B. M., Sawakuchi, H. O., Ward, N. D., Crump, B. C. 2017. Bacterial biogeography across the Amazon River-Ocean continuum. Frontiers in Microbiology, 8:882. doi: 10.3389/fmicb.2017.00882

Duarte, R. M., Smith, D. S., Val, A. L., Wood, C. M. 2016. Dissolved organic carbon from the upper Rio Negro protects zebrafish (*Danio rerio*) against ionoregulatory disturbances caused by low pH exposure. Scientific Reports, 6. doi:10.1038/srep20377

Dufrene, M., Legendre, P. 1997. Species assemblages and indicator species: The Need for a flexible asymmetrical approach. Ecological Monographs, 67:345–366. doi: 10.2307/2963459

Duperron, S., Halary, S., Habiballah, M., Gallet, A., Huet, H., Duval, C., Bernard, C., Benjamin, M. 2019. Response of fish gut microbiota to toxin-containing Cyanobacterial extracts: A microcosm study on the Medaka (*Oryzias latipes*). Environmental Science & Technology Letters, 6(6):341–347. doi: 10.1021/acs.estlett.9b00297

Audet-Gilbert, É., Sylvain, F-É., Bouslama, S., Derome, N. 2021. Microbiomes of clownfish and their symbiotic host anemone converge before their first physical contact. Microbiome, 9(1):109. doi: 10.1186/s40168-021-01058-1

Fadeev, E., Wietz, M., von Appen, W., Iversen, M.H., Nöthig, E., Engel, A., Grosse, J., Graeve, M., Boetius, A. 2021. Submesoscale physicochemical dynamics directly shape bacterioplankton community structure in space and time. Limnology and Oceanography. doi: 10.1002/lno.11799

Franke, M., Geier, B., Hammel, J. U., Dubilier, N., Leisch, N. 2021. Coming together-symbiont acquisition and early development in deep-sea bathymodioline mussels. Proceedings of the Royal Society: Biological sciences, 288(1957):20211044. doi: 10.1098/rspb.2021.1044

Gallet, A., Halary, S., Duval, C., Huet, H., Duperron, S., Marie, B. 2021. Disruption of fish gut microbiota composition and holobiont’s metabolome by cyanobacterial blooms. bioRxiv, 2021.09.08.459397. doi: 10.1101/2021.09.08.459397

Gaillardet, J., Dupre, B., Allegre, C. J., Negrel, P. 1997. Chemical and physical denudation in the Amazon River basin. Chemistry and Geology, 142:141–173. doi:10.1016/s0009-2541(97)00074-0

Ghosh, S., Whitley, C. S., Haribabu, B., Jala, V. R. 2021. Regulation of intestinal barrier function by microbial metabolites. Cellular and molecular gastroenterology and hepatology, 11(5):1463–1482. doi: 10.1016/j.jcmgh.2021.02.007

Gjessing, M. C., Spilsberg, B., Steinum, T. M., Amundsen, M., Austbo, L., Hansen, H., Colquhoun, D., Olsen, A. B. 2021. Multi-agent *in situ* hybridization confirms *Ca*. Branchiomonas cysticola as a major contributor in complex gill disease in Atlantic salmon. Fish and Shellfish Immunology Reports, 2. doi: 10.1016/j.fsirep.2021.100026

Goertz, S., de Menezes, A. B., Birtles, R. J., Fenn, J., Lowe, A. E., MacColl, A., Poulin, B., Young, S., Bradley, J. E., Taylor, C. H. 2019. Geographical location influences the composition of the gut microbiota in wild house mice (*Mus musculus domesticus*) at a fine spatial scale. PloS one, 14(9):e0222501. doi: 10.1371/journal.pone.0222501

Henderson, P. A., Crampton, W. G. R. 1997. A comparison of fish diversity and abundance between nutrient-rich and nutrient-poor lakes in the Upper Amazon. Journal of Tropical Ecology, 13:175–198. doi:10.1017/s0266467400010403

Henry, L.P., Bruijning, M., Forsberg, S.K.G., Ayroles, J. 2021. The microbiome extends host evolutionary potential. Nature Communication, 12:5141. doi: 10.1038/s41467-021-25315-x

Hermann, T. W., Stewart, D. J., Limburg, K. E., Castello, L. 2016. Unravelling the life history of Amazonian fishes through otolith microchemistry. Royal Society open science, 3(6):160206. doi: 10.1098/rsos.160206

Huson, D. H., Beier, S., Flade, I., Górska, A., El-Hadidi, M., Mitra, S., Ruscheweyh, H. J., Tappu, R. 2016. MEGAN community edition - interactive exploration and analysis of large-scale microbiome sequencing data. PLoS Computational Biology, 21;12(6):e1004957. doi: 10.1371/journal.pcbi.1004957

Junk, W.J., Soares, M.G., Carvalho, F.M. 1983. Distribution of fish species in a lake of the Amazon River floodplain near Manaus (Lago Camaleão), with special reference to extreme oxygen conditions. Amazoniana. 397– 431.

Krotman, Y., Yergaliyev, T.M., Alexander Shani, R., Avrahami, Y., Szitenberg, A. 2020. Dissecting the factors shaping fish skin microbiomes in a heterogeneous inland water system. Microbiome, 8(9). doi: 10.1186/s40168-020-0784-5

Lavoie, C., Wellband, K., Perreault, A., Bernatchez, L., Derome, N. 2021. Artificial rearing of Atlantic salmon juveniles for supportive breeding programs induces long-term effects on gut microbiota after stocking. Microorganisms, 9:1932. doi: 10.3390/microorganisms9091932

Legendre, P., Anderson, M.J. 1999. Distance-based redundancy analysis: Testing multispecies responses in multifactorial ecological experiments. Ecological Monographs, 69:1–24. doi: 10.1890/0012-9615(1999)069[0001:DBRATM]2.0.CO;2

Legrand, T., Catalano, S. R., Wos-Oxley, M. L., Stephens, F., Landos, M., Bansemer, M. S., Stone, D., Qin, J. G., Oxley, A. 2018. The inner workings of the outer surface: Skin and gill microbiota as indicators of changing gut health in yellowtail kingfish. Frontiers in microbiology, 8:2664. doi: 10.3389/fmicb.2017.02664

Leroux, N., Sylvain, F-É, Normandeau, E, Holland, A., Val, A.L., Derome, N. 2022. Evolution of an Amazonian fish is driven by allopatric divergence rather than ecological divergence. Frontiers in Ecology and Evolution, 10:875961. doi: 10.3389/fevo.2022.875961

Lim, W. G., Tong, T., Chew, J. 2020. *Chryseobacterium indologenes* and *Chryseobacterium gleum* interact and multiply intracellularly in *Acanthamoeba castellanii*. Experimental parasitology, 211:107862. doi: 10.1016/j.exppara.2020.107862

Llewellyn, M. S., Boutin, S., Hoseinifar, S. H., Derome, N. 2014. Teleost microbiomes: the state of the art in their characterization, manipulation and importance in aquaculture and fisheries. Frontiers in microbiology, 5:207. doi: 10.3389/fmicb.2014.00207

Llewellyn, M.S., McGinnity, P., Dionne, M., Létourneau, J., Thonier, F., Carvalho, G.R., Creer, S., Derome, N. 2016. The biogeography of the Atlantic salmon (*Salmo salar*) gut microbiome. The ISME Journal, 10:1280–1284. doi: 10.1038/ismej.2015.189

Lindh, M. V., Pinhassi, J. 2018. Sensitivity of bacterioplankton to environmental disturbance: A review of Baltic Sea field studies and experiments. Frontiers in Marine Science, 5:361. doi: 10.3389/fmars.2018.00361

Margulis, L., Fester, R. 1991. Symbiosis as a Source of Evolutionary Innovation: Speciation and Morphogenesis. MIT Press, Cambridge, Mass. and London.

McDevitt-Irwin, J. M., Baum, J. K., Garren, M., Vega Thurber, R. L. 2017. Responses of coral-associated bacterial communities to local and global stressors. Frontiers in Marine Science, 4:262. doi: 10.3389/fmars.2017.00262

McMurdie, P. J., S. Holmes. 2013. phyloseq: An R package for reproducible interactive analysis and graphics of microbiome census data. Plos One, 8. doi: 10.1371/journal.pone.0061217

Menke, S., Gillingham, M. A. F., Wilhelm, K., Sommer, S. 2017. Home-made cost effective preservation buffer is a better alternative to commercial preservation methods for microbiome research. Frontiers in Microbiology, 8. doi: 10.3389/fmicb.2017.00102

Minich, J. J., Petrus, S., Michael, J. D., Michael, T. P., Knight, R., Allen, E. E. 2020. Temporal, environmental, and biological drivers of the mucosal microbiome in a wild marine fish, Scomber japonicus. mSphere, 5(3):e00401–20. doi: 10.1128/mSphere.00401-20

Mitchell, S. O., Steinum, T. M., Toenshoff, E. R., Kvellestad, A., Falk, K., Horn, M., Colquhoun, D. J. 2013. *’Candidatus Branchiomonas cysticola’* is a common agent of epitheliocysts in seawater-farmed Atlantic salmon *Salmo salar* in Norway and Ireland. Diseases of aquatic organisms, 103(1):35–43. doi: 10.3354/dao02563

Morris, C., Val, A. L., Brauner, C. J., Wood, C. M. 2021. The physiology of fish in acidic waters rich in dissolved organic carbon, with specific reference to the Amazon basin: Ionoregulation, acid-base regulation, ammonia excretion, and metal toxicity. Journal of experimental zoology, Part A, Ecological and integrative physiology, 335(9-10):843– 863. doi: 10.1002/jez.2468

Nossa, C. W., Oberdorf, W. E., Yang, L., Aas, J. A., Paster, B. J., Desantis, T. Z., Brodie, E. L., Malamud, D., Poles, M. A., Pei, Z. 2010. Design of 16S rRNA gene primers for 454 pyrosequencing of the human foregut microbiome. World Journal of Gastroenterology, 16:4135–4144. doi:10.3748/wjg.v16.i33.4135

Jari Oksanen, F., Blanchet, G., Friendly, M., Kindt, R., Legendre, P., McGlinn, D., Minchin, P. R., O’Hara, R. B., Simpson, G. L., Solymos, P., Henry, M., Stevens, H., Szoecs, E., Wagner, H. 2020. vegan: Community ecology package. R package version 2.5–7. https://CRAN.R-project.org/package=vegan

Pang, X., Shen, H., Niu, Y., Sun, X., Chen, J., Xie, P. 2014. Dissolved organic carbon and relationship with bacterioplankton community composition in 3 lake regions of Lake Taihu, China. Canadian journal of microbiology, 60(10):669–680. doi: 10.1139/cjm-2013-0847

Petry, P., Bayley, P. B., Markle, D. F. 2003. Relationships between fish assemblages, macrophytes and environmental gradients in the Amazon River floodplain. Journal of fish biology, 63: 547–579. doi:10.1046/j.1095-8649.2003.00169.x

Pires, T., Borghezan, E. A., Machado, V. N., Powell, D. L., Röpke, C. P., Oliveira, C., Zuanon, J., Farias, I. P. 2018. Testing Wallace’s intuition: water type, reproductive isolation and divergence in an Amazonian fish. Journal of evolutionary biology, 31(6):882–892. doi: 10.1111/jeb.13272

Pratte, Z. A., Besson, M., Hollman, R. D., Stewart, F. J. 2018. The gills of reef fish support a distinct microbiome influenced by host-specific factors. Applied and environmental microbiology, 84(9):e00063–18. doi: 10.1128/AEM.00063-18

Riiser, E. S., Haverkamp, T. H. A., Varadharajan, S., Borgan, O., Jakobsen, K. S., Jentoft, S., Star, B. 2020. Metagenomic shotgun analyses reveal complex patterns of intra- and interspecific variation in the intestinal microbiomes of codfishes. Applied and Environmental Microbiology, 86(6). doi:10.1128/aem.02788-19

Rodriguez, M. A., Lewis, W. M. 1997. Structure of fish assemblages along environmental gradients in floodplain lakes of the Orinoco River. Ecological Monographs. 67:109–128, doi:10.2307/2963507

Saint-Paul, U., ZUanon, J., Correa, M. A. V., Garcia, M., Fabre, N. N., Berger, U., Junk, W. J. 2000. Fish communities in central Amazonian white- and blackwater floodplains. Environmental Biology of Fishes, 57:235–250. doi:10.1023/a:1007699130333

Santos-Júnior, C. D., Kishi, L. T., Toyama, D., Soares-Costa, A., Oliveira, T. C., de Miranda, F. P., Henrique-Silva, F. 2017. Metagenome sequencing of Prokaryotic microbiota collected from rivers in the Upper Amazon Basin. Genome announcements, 5(2) :e01450–16. doi : 10.1128/genomeA.01450-16

Satinsky, B. M., Fortunato, C. S., Doherty, M., Smith, C. B., Sharma, S., Ward, N. D., Krusche, A. V., Yager, P. L., Richey, J. E., Moran, M. A., Crump, B. C. 2015. Metagenomic and metatranscriptomic inventories of the lower Amazon River, May 2011. Microbiome, 3:39. doi: 10.1186/s40168-015-0099-0

Satinsky, B. M, Crump, B. C., Smith, C. B., Sharma, S., Zielinski, B. L., Doherty, M., Meng, J., Sun, S., Medeiros, P. M., Paul, J. H., Coles, V. J., Yager, P. L., Moran, M. A. 2014. Microspatial gene expression patterns in the Amazon River Plume. Proceedings of the National Academy of Science of the United-States of America, 111(30):11085– 90. doi: 10.1073/pnas.1402782111

Satinsky BM, Zielinski BL, Doherty M, Smith CB, Sharma S, Paul JH, Crump, B. C., Moran, M. A. 2014. The Amazon continuum dataset: quantitative metagenomic and metatranscriptomic inventories of the Amazon River plume, June 2010. Microbiome, 2:17. doi: 10.1186/2049-2618-2-17.

Sevellec, M., Derome, N., Bernatchez, L. 2018. Holobionts and ecological speciation: the intestinal microbiota of lake whitefish species pairs. Microbiome, 6. doi:10.1186/s40168-018-0427-2

Smith, C. C. R., Snowberg, L. K., Caporaso, J. G., Knight, R., Bolnick, D. I. 2015. Dietary input of microbes and host genetic variation shape among-population differences in stickleback gut microbiota. Isme Journal, 9(11) :2515–2526. doi:10.1038/ismej.2015.64

Sioli, H. 1984. The Amazon limnology and landscape ecology of a mighty tropical river and its Basin. Dr. Junk Publisher, Dordrecht.

Stevens, E. J., Bates, K. A., King, K. C. 2021. Host microbiota can facilitate pathogen infection. PLoS pathogens, 17(5):e1009514. doi: 10.1371/journal.ppat.1009514

Sylvain, F-É., Holland, A., Audet-Gilbert, É., Val, A. L., Derome, N. 2019. Amazon fish bacterial communities show structural convergence along widespread hydrochemical gradients. Molecular Ecology, 28:3612–3626. doi: 10.1111/mec.15184

Sylvain, F. É., Holland, A., Bouslama, S., Audet-Gilbert, É., Lavoie, C., Val, A. L., Derome, N. 2020. Fish skin and gut microbiomes show contrasting signatures of host species and habitat. Applied and environmental microbiology, 86(16):e00789–20. doi: 10.1128/AEM.00789-20

Sylvain, F-É., Bouslama, S., Holland, A., Leroux, N., Mercier, P., Val, A.L., Derome, N. 2021. The Amazon River microbiome, a story of humic carbon. bioRxiv. doi: 10.1101/2021.07.21.453257

Sylvain, F-É., Cheaib, B., Llewellyn, M., Correia, T. G., Fagundes, D. B., Val, A. L., Derome, N. 2016. pH drop impacts differentially skin and gut microbiota of the Amazonian fish tambaqui (*Colossoma macropomum*). Scientific Reports, 6:32032. doi: 10.1038/srep32032

Sylvain, F-É., Derome, N. 2017. Vertically and horizontally transmitted microbial symbionts shape the gut microbiota ontogenesis of a skin-mucus feeding discus fish progeny. Scientific Reports, 7:5263. doi: 10.1038/s41598-017-05662-w

Sylvain, F-É., Normandeau, E., Holland, A., Val, A. L., Derome, N. 2021. Genomics of Serrasalmidae teleosts through the lens of microbiome fingerprinting. Authorea. doi: 10.22541/au.163256091.18930269/v1

Pires, T. H. S., Campos, D. F., Röpke, C. P., Sodré, J., Amadio, S., Zuanon, J. 2015. Ecology and life-history of *Mesonauta festivus*: biological traits of a broad ranged and abundant Neotropical cichlid. Environmental biology of fishes, 98:789–799. doi: 10.1007/s10641-014-0314-z

Toyama, D., Kishi, L. T., Santos-Júnior, C. D., Soares-Costa, A., de Oliveira, T. C., de Miranda, F. P., Henrique-Silva, F. 2016. Metagenomics analysis of microorganisms in freshwater lakes of the Amazon Basin. Genome announcements, 4(6):e01440–16. doi: 10.1128/genomeA.01440-16

Ulluwishewa, D., Anderson, R. C., McNabb, W. C., Moughan, P. J., Wells, J. M., Roy, N. C. 2011. Regulation of tight junction permeability by intestinal bacteria and dietary components. The Journal of nutrition, 141(5):769–776. doi: 10.3945/jn.110.135657

Val, A. L., Almeida-Val, V. M. F. 1995. Fishes of the Amazon and their environment. Springer-Verlag Berlin Heidelberg.

Webster, T. M. U., Consuegra, S., Hitchings, M., de Leaniz, C. G. 2018. Interpopulation variation in the Atlantic salmon microbiome reflects environmental and genetic diversity. Applied and Environmental Microbiology, 84(16). doi: 10.1128/aem.00691-18

